# EcoLiDAR: An economical LiDAR scanner for ecological research

**DOI:** 10.1101/2024.01.31.578255

**Authors:** Calebe Pereira Mendes, Norman T-Lon Lim

## Abstract

Despite recent popularization and widespread use in modern electronic devices, LiDAR technology remains expensive for research purposes, in part due to the very high performance offered by commercially available LiDAR scanners. However, such high performance is not always needed, and the expensive price ends up making LiDAR scanners inaccessible for research projects with reduced budget, such as those in developing countries. Here we designed and built a simple ground-based LiDAR scanner, with performance sufficient to fulfil the requirements for a variety of ecological research projects, while being cheap and easy to build. We managed to assemble a LiDAR scanner under 400 USD (as of 2021), and it is simple enough to be built by personnel with minimal engineering background. We also demonstrated the quality of the resulting point clouds by scanning a test site and producing some common LiDAR products. Although not adequate for mapping large area due to its limited range, our LiDAR design is open, customizable, and can produce adequate results while costing ∼1% of “low-cost” scanners available in the market. As such, our LiDAR scanner opens a world of new opportunities, particularly for projects in developing countries.

## Introduction

In the last 20 years, LiDAR (light detection and ranging) technology matured to a point where even household and personal devices, such as smartphones and robot vacuum cleaners, are equipped with LiDAR sensors. By measuring the time-of-flight of light emitted by a laser diode (or by a focused LED in simpler sensors), LiDAR sensors can precisely measure distance to the illuminated target [1], and thus is an excellent sensor option for navigation systems and for three-dimensional object recognition [2, 3]. As a research tool, LiDAR technology has been used to investigate a variety of subjects, from creating large-scale digital elevation models [2, 4], to mapping the three-dimensional structure of forests [5–7], and measuring the morphometric dimensions of living animals [8]. To achieve such a variety of applications, LiDAR devices can be divided into five deployment methods [5]: 1) spaceflight LiDAR; 2) airborne LiDAR from manned aircraft; 3) airborne LiDAR from unmanned aerial vehicle; 4) mobile ground-based LiDAR; and 5) static ground-based LiDAR.

When operated from fast-moving platforms, such as satellites or aircrafts, LiDAR sensors can quickly scan very large extensions of land to produce digital elevation models and land use maps, and to measure vegetation coverage in large landscapes [7, 9]. However, due to the dispersion of the laser beam, farther measurement distances also result in larger laser footprints (i.e., area illuminated by a single laser bean), and consequently, the point density of the scanned surfaces is reduced [1, 5]. As a result, the final products of large footprint LiDAR devices are similar in dimension and resolution to those produced by other remote sensing techniques, such as radar and high-resolution photographic imagery [10]. Indeed, the similar resolution allows satellite and airborne LiDAR products to be used in combination with other remote sensing techniques to generate improved products, such as high quality forest biomass estimations from combining LiDAR and synthetic-aperture radar [11]. On the other hand, LiDAR devices that operate near the illuminated targets, such as from low-flying unmanned aerial vehicles and ground-based LiDAR scanners, can scan small areas with a small laser footprint and thus generate extremely high point densities [5]. In turn, LiDAR products with high point density allow precise mapping of fine-scale three-dimensional structures, such as canopy leaf distribution [12], leaf orientation [13], shape and volume of trunks and branches [14], and even the identification of the tree species [15].

Despite the usefulness and large-scale adoption of LiDAR technology in both research and commercial applications, it is still a relatively expensive technology, with systems costing up to 40,000 USD still being considered as “low-cost” [16]. Some of the cheapest options for static ground-based LiDAR scanners currently available in the market include the Leica BLK360, with prices around 19,000 USD [17], and the FARO Focus M 70, costing around 25,000 USD [18]. Although within the purchase power of well-funded research institutions and laboratories, such price tags can be prohibitive for research teams with lower budgets. To put things into perspective, for the price of a “low-cost” Leica BLK360 LiDAR device, an U.S.-based laboratory can hire a postdoctoral fellow for five months [19]. This situation becomes more problematic for research institutions located in developing countries, where the same Leica BLK360 scanner would be equivalent to 22 months of salary for a postdoc in Brazil (based on official wages of the National Council for Scientific and Technological Development (CNPq 2022) and a ratio of 1 BRL to 0.21 USD [20]). To further complicate matters, most of earth’s biodiversity hotspots are located in developing countries [21], where research institutions in those countries have easy access to and extensive expertise on the local environment, but may not have access to the expensive equipment needed to better study these vulnerable ecosystems (e.g., tropical rainforests with complex three-dimensional vegetation).

In a similar way that open-source software democratised analytic tools previously locked behind expensive licences [22], the democratisation of LiDAR technology and its usage by research institutions with limited funds could play a crucial role on boosting the development of environmental science. Although high-performance electronics are generally expensive, a low-budget LiDAR device is generally sufficient to produce usable data for many applications, such as mapping small vegetation plots. Moreover, simple LiDAR devices can be easily manufactured from cheap and available electronic components. Therefore, the objective of this manuscript is to provide an open-source design to a simple, cheap, and functional LiDAR scanner (named as EcoLiDAR), such that the building of this device can be accomplished by someone with minimal engineering background (although basic electronic knowledge still required). Additionally, we showcase the data output to demonstrate its potential and limitations.

We opted for a stationary ground-based LiDAR scanner, as opposed to a mobile terrestrial or airborne LiDAR scanner, to keep the equipment as simple as possible. Mobile LiDAR scanners are intrinsically more complex than the stationary options, requiring very high scanning speeds and relying on precise orientation sensors [1], while keeping a reduced size and weight. These requirements have a direct impact on the build complexity and the final price tag.

The following characteristics were also defined as requirements for the system: 1) the LiDAR scanner can be attached to a tripod to produce an accurate 3D scan of the surrounding environment, scan objects up to 30 m away, within < 30 minutes scan time, while operating with off-grid power. 2) It must be simple enough to be built by personnel with limited engineering background. 3) It should be as cheap as possible to be a viable option for small-budget research projects and/or for researchers from countries with weaker currencies. 4) The parts must be widely available in the market and the required software for its operation must be available for free.

## Materials and methods

### Laser rangefinder

As the main sensor of a LiDAR device, the specifications of the laser rangefinder define the final capabilities of the device. With this in mind, we chose the Garmin LIDAR-Lite v3HP [23] as the laser rangefinder sensor, mainly due to its advertised 40-m range, capability to operate at outdoor environments, and low price; the outdoor capability and 40-m range are features not commonly found in other similar-priced laser rangefinders. Another crucial point of consideration was the I^2^C communication protocol and the large availability of online libraries (both in Python and C++) for interfacing the LIDAR-Lite v3HP with microcontrollers. On the other hand, the main drawback of the LIDAR-Lite v3HP is its slow sampling rate, advertised as >1 kHz but only achievable under controlled indoor situations. For outdoor conditions, ∼300 Hz appears to be the maximum functional sampling rate. Other drawbacks that must be considered are the low resolution (1 cm) and low accuracy (2.5 cm) of the distance measurements, as well as the high divergence of the laser (8 mRad), which increases the laser footprint as distance increases [23]. These laser-related drawbacks are also found in other laser rangefinders in the same price range.

### Processing unit

Working as the equipment’s brain, the processing unit is responsible for running the scanner’s software, controlling the sensors and actuators, and saving the scanned data for subsequent analyses. The choice of a processing unit also defines the programming language to be used on the scanner’s software. After some initial experimentation between an Arduino uno Rev3 [24], Raspberry Pico [25], and Raspberry Pi Zero [26], we chose the latter as the LiDAR’s processing unit. The decision for the Raspberry Pi Zero was made due to the built-in SD card reader and Wi-Fi capability, and the vast availability of software and documentation since it can run operation systems, such as the Raspberry Pi OS and other Linux distributions [27]. In fact, the Raspberry Pi Zero is better described as a single-board computer, while the Arduino uno Rev3 and Raspberry Pico are typical microcontrollers [24, 25]. It is noteworthy that the higher processing speed of the Raspberry Pi Zero is not necessarily an improvement to the LiDAR scanner, since both microcontrollers have sufficient performance to run the LiDAR software at a 300 Hz sampling rate. Moreover, the use of an operation system in the Raspberry Pi Zero has the drawback of increasing the LiDAR booting time. However, the higher performance of the Raspberry Pi Zero can be potentially useful for future-proofing, such as changing the LIDAR-Lite v3HP for another laser rangefinder with a higher sampling rate.

### Two-axis gimbal

The Garmin LIDAR-Lite v3HP, as with other laser rangefinders, does not have any moving parts and can only measure distances within a single laser beam (i.e., one dimension) emitted by the sensor [23]. Therefore, the laser rangefinder needs to be attached to a laser deflection mechanism to allow a 360° scan of the surrounding area [28]. Alternatively, the laser rangefinder can be attached to a two-axis gimbal, which provides the pan-and-tilt movements needed for a 360° scan. To better accommodate the electronic components, we designed the gimbal from scratch using a mixture of commercially available and 3D printed parts. The two-axis gimbal can be separated in two sections, each responsible for one axis of rotation (i.e., pan and tilt) and containing one stepper motor (Fig 1).

**Fig 1.**
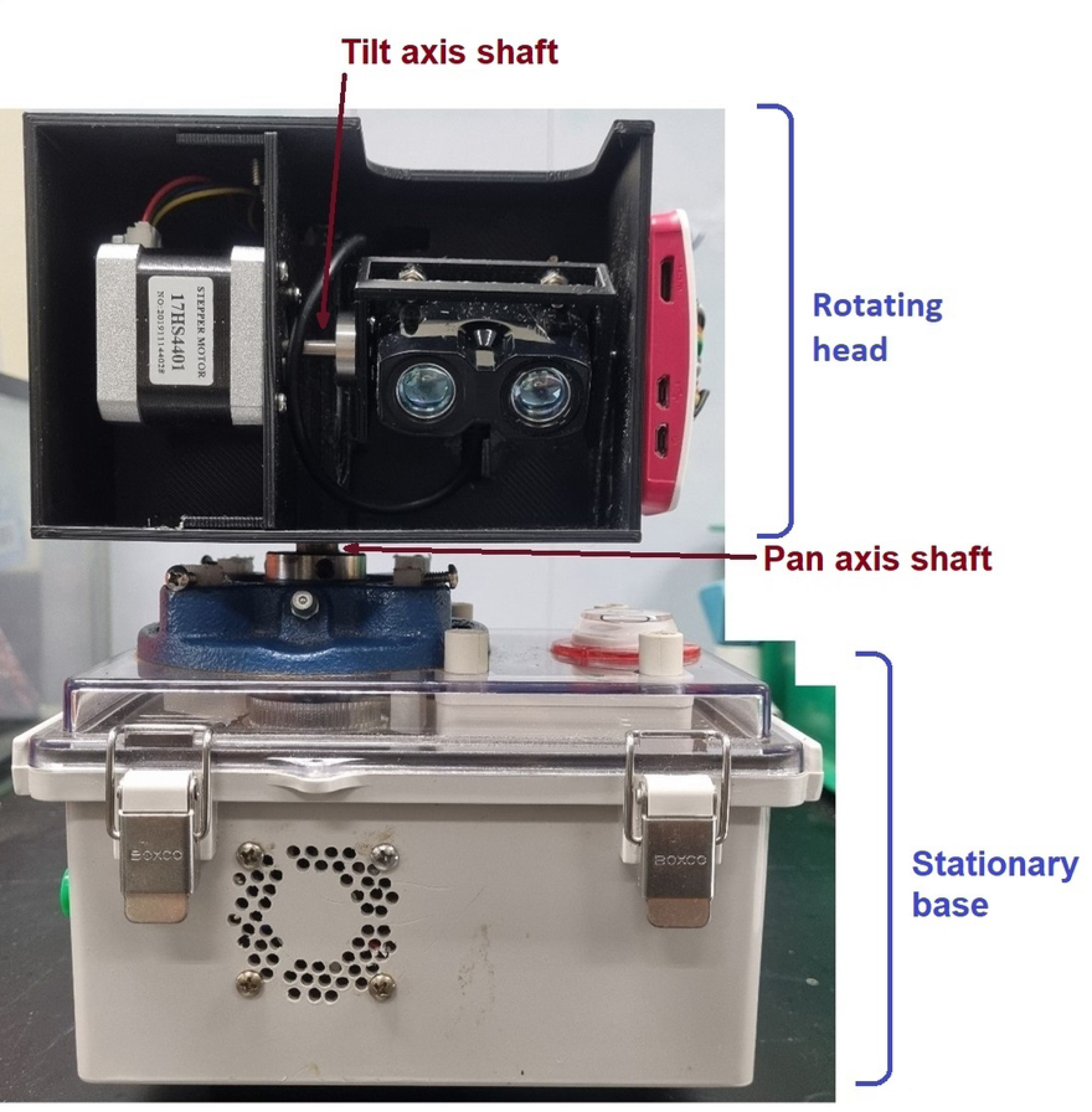
EcoLiDAR scanner. The two axes of rotation are pointed by red arrows. The division between the “rotating head” and “stationary base” is also displayed.

The pan-axis motion of the gimbal also separates the device in two portions: the “rotating head”, which rotates, and the “stationary base”, which is stationary and connects to a tripod during operation (Fig 1). The pan-axis motion is created by a flanged bearing bolted to the stationary base and connected to a vertical steel shaft, 12 mm diameter and 120 mm long. The vertical shaft, which can rotate due to the bearing, is firmly connected on its upper end to the “rotating head”, which is a 3D-printed case containing the laser rangefinder, the processing unit, and the gimbal section responsible for the tilt motion (Fig 2). The rotating head is 145 × 105 × 100 mm and was 3D-printed in PETG. However, in the lack of a 3D-printer, the same case can be build using proper woodworking techniques or from plastic sheets using plastic welding.

**Fig 2.**
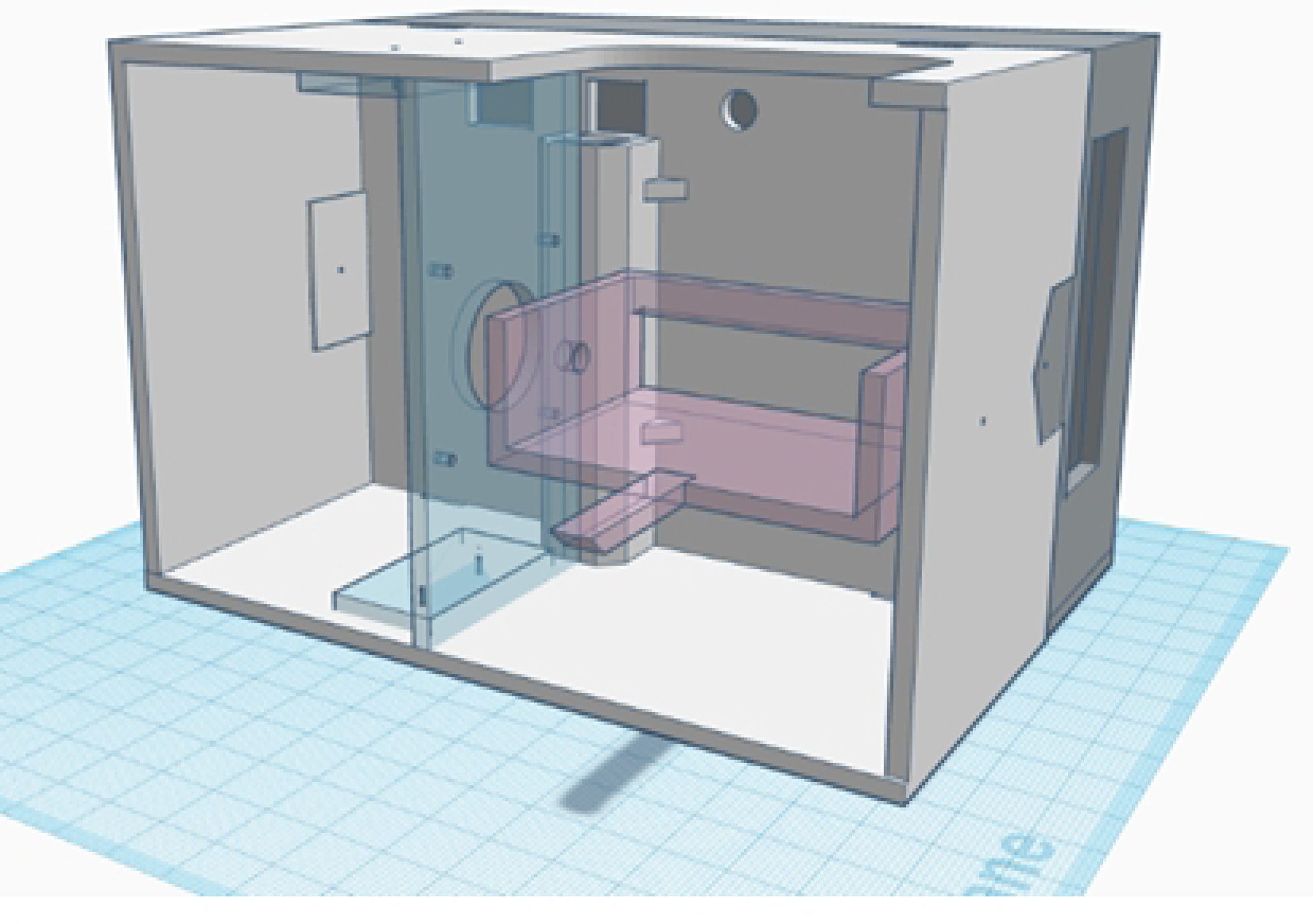
Rotating head. The “rotating head” contains the rotating components of the LiDAR scanner. The head can be divided in the main box (white), the electronic lid (gray), the stepper motor support (blue) and LiDAR support (pink). The 3D printing files are available in a GitHub repository.

The lower end of the vertical shaft, inside the stationary base, is connected to a timing gear, which is powered by a stepper motor through a GT2 timing belt, in a 3:1 gear ratio (S1 Fig). The electric connection between the rotating head and the stationary base is made by eight wires running inside the vertical shaft, while a slip ring located in the bottom of the shaft allows the wires to rotate freely. As the slip ring can cause interference to high frequency communication, the processing unit was placed in the rotating head instead of the stationary base. Finally, the tilt motion is created by directly attaching the laser rangefinder to a stepper motor that is located in the rotating head. It is important to highlight that the laser rangefinder is not located in the centre of the vertical axis due to the weight distribution of the rotating head. This design helps to avoid vibrations during operation, but causes an offset in the collected data, which need to be corrected by software. An R script is provided in the supplementary materials to correct the offset (S2 Code).

### Stepper motors

For powering the two-axis gimbal, a pair of Nema 17 stepper motors (MotionKing, Changzhou, model 17HS4401) were used [29]. The two-phase hybrid stepper motor has 200 steps and 40 N.cm of torque. Subsequent tests demonstrated that weaker motors (∼20 N.cm) would suffice, while being lighter and cheaper. The motors were operated at 30 V, with the current limited to 1.5 A by a A4988 stepper motor driver module [30]. The A4988 driver was chosen due to its high availability in the market, low price, ease of use, and it being interchangeable with other driver modules, such as the DRV8825, MP6500 and DRV8880 (although minor software adjustments may be required when swapping between different driver models). The A4988 also allows for micro-stepping up to 1/16th of step, which is important for controlling the resolution of the LiDAR scan.

The rotating head spinning speed is defined by:

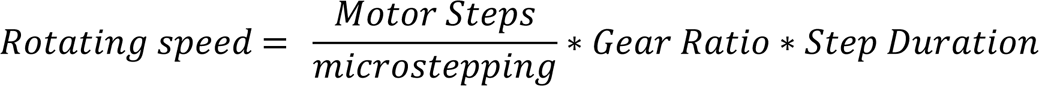

where the number motor steps is defined by the motor design (200 steps for the model 17HS4401), the microstepping is defined by the stepper motor driver and can range from 1 (i.e., fullstep) to 1/16 when using the A4988, the gear ratio is 3:1 (as described in the 2-axis gimbal section), and the step duration is the time interval between sequential steps (or microsteps) when the motor is operating. The step duration can be defined by software but is limited to a minimum interval of 3.3 milliseconds, to not exceed the 300 Hz maximum sampling rate of the laser rangefinder. Therefore, when operating at the maximum sampling rate (300 Hz) and using a 1/4 microstepping, each revolution of the rotating head takes 7.92 seconds and produces 2400 distance measurements (0.15° degrees apart from each other, every 3.3 milliseconds).

The total duration of a scan depends on the duration of the rotating head revolutions multiplied by the number of revolutions in a complete scan. In turn, the number of revolutions required to complete a scan is defined by:

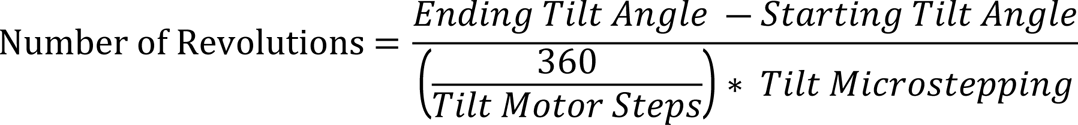

where the starting and ending tilt angles are the vertical angles in which the laser rangefinder will start and end the scan (considering a horizontal coordinate system, with zenith = 90°, horizon = 0° and nadir = −90°). Note that the number of tilt motor steps is defined by the motor design (200 steps for the model 17HS4401), while the tilt microstepping is controlled by the stepper motor driver. Therefore, a scan starting with the laser rangefinder pointing at −40° and finishing at 90° (zenith), and with a 1/4 tilt-axis microstepping (using the 17HS4401 stepper motor) will contain 289 revolutions. Considering a revolution duration of 7.92 seconds, the complete scan duration is roughly 38 minutes, producing a point cloud with 693,600 points. In this design, the tilt-axis microstepping is defined by hardware and requires some wire soldering in order to be changed; however, software control of the tilt-axis microsteps can be easily implemented by adding wires connecting the processing unity to the tilt-axis motor driver and by simple changes in the LiDAR’s software.

### Battery and voltage regulators

To allow the LiDAR scanner to operate off-grid, a rechargeable 12 V, 7 Ah sealed lead-acid battery was used as a power supply. The battery is connected to the scanner by a two-core power cable with a pair of alligator clips at one end (for connection to the battery terminals) and a male 5.5 mm DC jack on the other end (to connect with a female 5.5 mm DC jack at the scanner’s stationary base). It is important to highlight that each lead-acid battery weighs ∼2 kg, which is a considerable drawback for fieldwork logistics. The use of lithium batteries may be a good alternative for reducing the weight.

Since different components of the LiDAR system operate at different voltage levels, three voltage regulators were used. One LM2596 adjustable step-down buck converter was used to reduce the voltage from 12 V, supplied by the battery, to the 5 V required by the Raspberry Pi Zero, LiDAR module, motor drivers, and cooling fan. Meanwhile, two XL6009 adjustable step-up buck converters were used to increase the voltage from 12 V, supplied by the battery, to the 30 V required for a reliable operation of the stepper motors. To avoid overheating the XL6009 step-up converters, each XL6009 supplies current for a single stepper motor only.

### Stationary base

To accommodate all the components and to maintain structural integrity of the system, the stationary base was built from a 20 × 15 × 10 cm electrical junction box. At one of the box sides, a female 5.5 mm DC jack was added to connect the battery power supply. Two push buttons (“start” and “stop”) and a power switch were also added (Fig 3). Two openings were made to allow air cooling, and a 5 V 40 × 40 × 10 mm fan was added to produce air flow. Finally, a tripod quick-release plate was firmly attached by screws to the bottom of the box, near its center of mass, allowing the stationary base to be directly connected to a tripod head.

**Fig 3.**
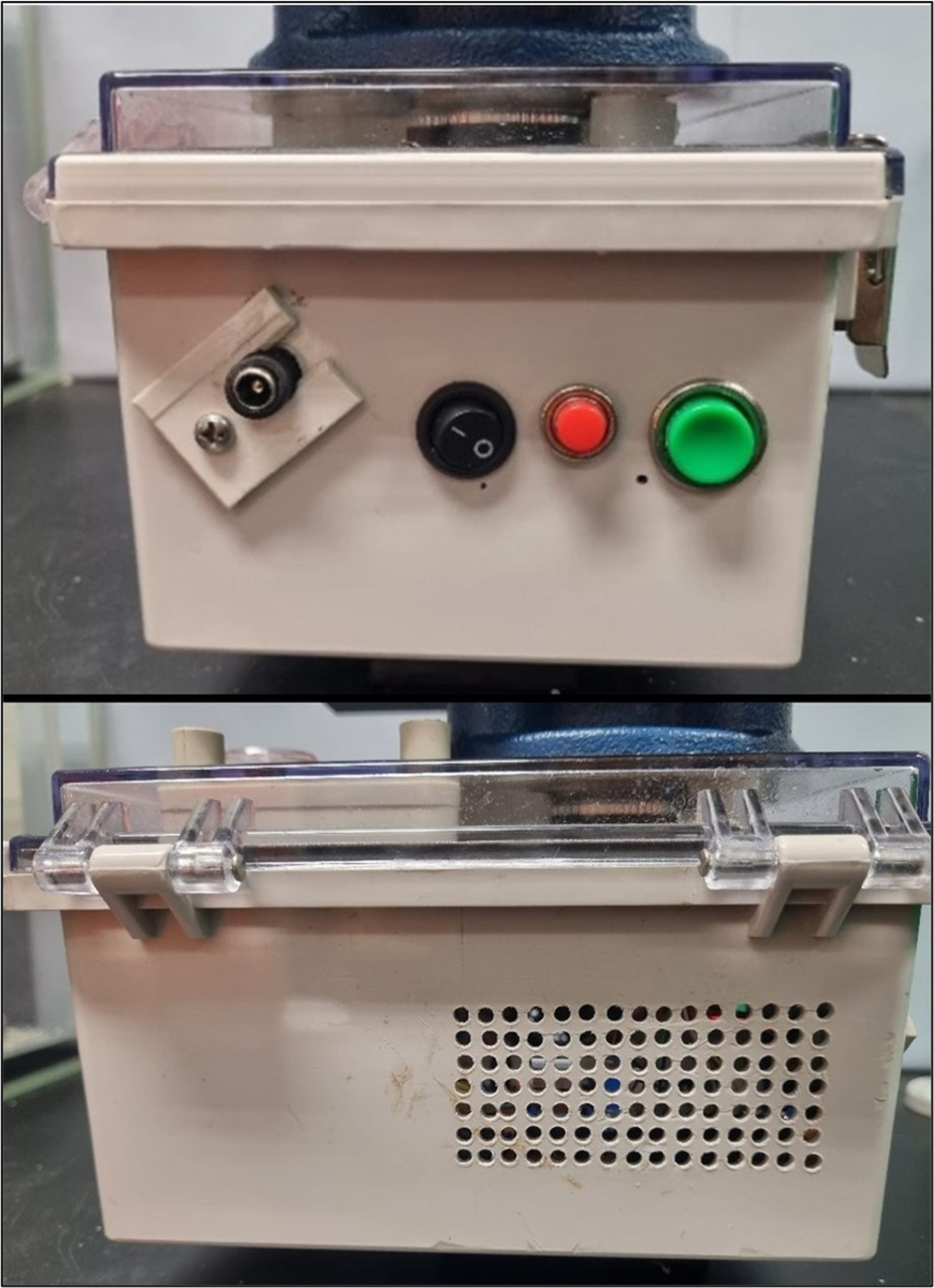
Stationary base. The “stationary base” contains the non-rotating components of the LiDAR scanner. The box smaller side (top) holds the start button in green, the stop button in red, the power switch and a 5.5 mm DC jack. The two long sides have cooling vents (bottom), with the air flow made by a single 5V fan.

### Wiring

The electric circuit of the LiDAR scanner was assembled on two solderable perforated breadboards, one for the stationary base and one for the rotating head, following the general schematics show in the Figure 4. Note that Figure 4 did not include the cooling fan, which is directly wired to the 5 V provided by the LM2577 step-down module.

**Fig 4.**
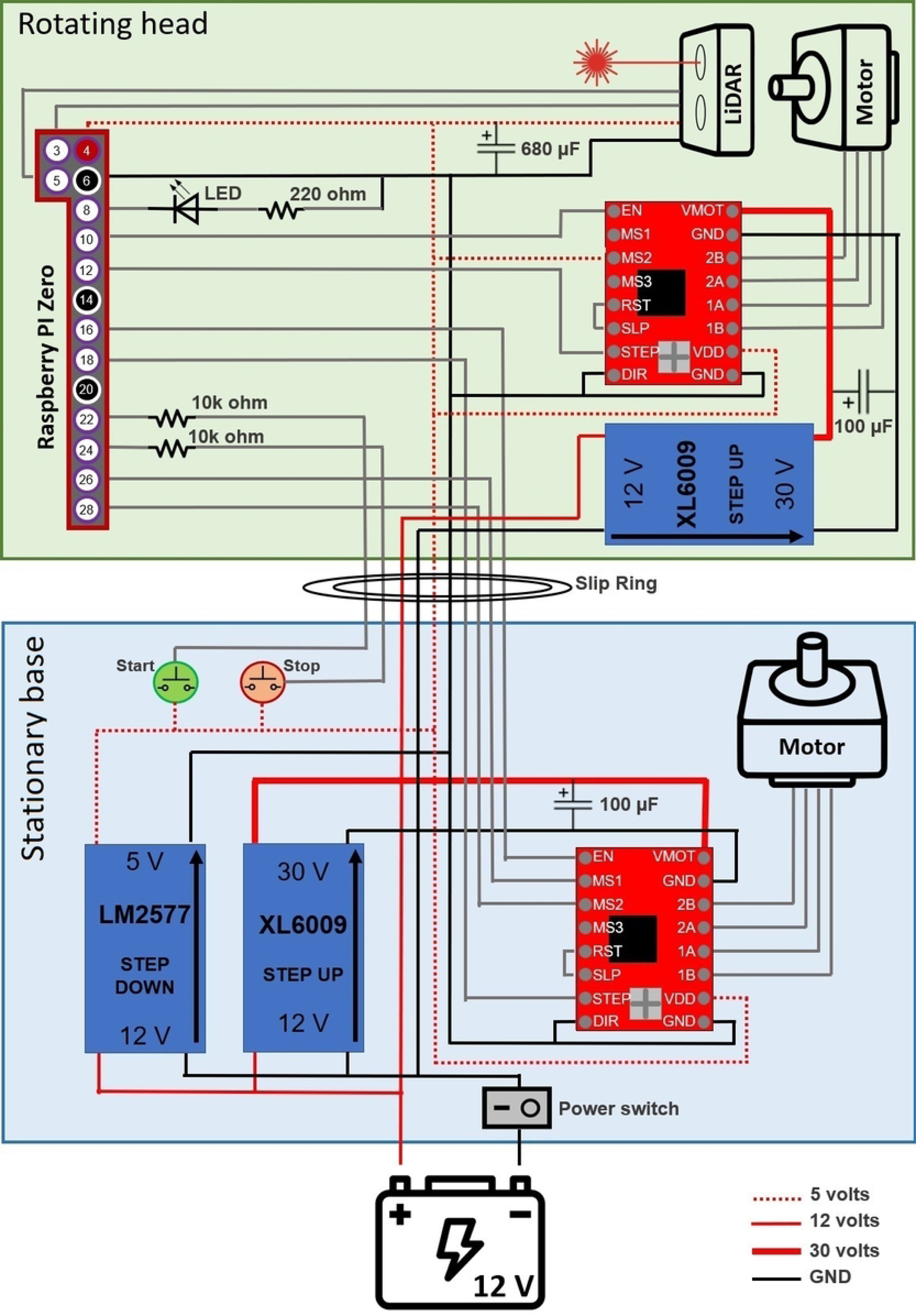
Diagram of the main circuit of the EcoLiDAR scanner. The image shows the Raspberry pin numbers, and not the General-Purpose Input-Output (GPIO) numbers. The wiring of the cooling fan is not displayed.

### Software

The software setup consists of the installation of the Raspberry Pi operating system (version 10; Buster) and the enabling the I^2^C interface, which comes turned off by default in the Raspberry Pi operating system. To control the LiDAR functions, instructions were written in Python 3 (see S3 Code) and saved as a .py file, which was set to be automatically executed on boot using the Linux task scheduler utility “Crontab”. Finally, a folder was created to save the scan files, which consist of a .txt file containing a single table with one LiDAR point per line, as well as its coordinates (in a horizontal coordinate system) and the time of measurement in milliseconds from the start of the scan. The retrieval of the scanned data can be performed via connecting the scanner’s Raspberry Pi Zero to a Wi-Fi network and downloading the files to an external device through a remote desktop software. More details about how to access the LiDAR’s files are described in the detailed user’s manual (S4 Manual).

### Orientation

To convert the laser rangefinder measurements into a 3D point cloud, the precise orientation of the laser during each measurement needs to be recorded. Since the motion of stepper motors can be tracked with high precision for extended periods of time, no additional sensors are required, provided that the laser orientation in the start of the scan is known. By recording the sequence of angular movements performed by the stepper motors during the scan, it is possible to calculate the precise orientation of the laser at any moment by backtracking the motor movements from the initial laser orientation. For instance, if a scan starts with the laser pointed to the north (bearing = 0°) and a tilt angle pointing to the horizon (tilt angle = 0°), the pan-axis motor (turning clockwise, with a 1/4 microstepping and a 3:1 gear ratio, as described above in the “Stepper motors” section) will move the laser in the pan axis by 0.15° per step (or −0.15° in the case of anti-clockwise motion). The new laser orientation after one step is therefore bearing = 0.15°, tilt angle = 0°. The same logic applies to the tilt-axis motion. Note that the bearing is a circular variable (therefore 0° = 360°). Finally, taking advantage of the fact that stepper motors did not accumulate angular imprecisions during operation, the precision of the orientation is maintained along the entire scan. However, step skipping (i.e., when the motor shaft does not move in synchrony with the coils) must be avoided at all costs, since it causes a cumulative error and affects the angular precision of all subsequent measurements.

### Laser alignment

The precise alignment of the laser rangefinder within the two-axis gimbal is crucial to ensure that the laser’s orientation during scanning aligns accurately with the angles recorded by the software. The software calculates the laser’s orientation by backtracking the gimbal motion from a predefined starting position. For instance, if the software assumes a scan to commence at bearing 0° (north) and a tilt angle of 40° below the horizon, it is vital to have the laser rangefinder precisely directed in the same direction at the start of the scan. To achieve this, laser alignment can be performed in a dark room utilizing a camera capable of detecting the infrared (IR) laser emitted by the rangefinder. Most low-light cameras, including webcams and smartphones, can detect the wavelengths emitted by LiDAR modules, enabling visual measurement of the laser’s position (S5 Fig).

The alignment process involves placing the rotating head in a fixed position while disabling the gimbal stepper motors through software or hardware (e.g., disconnecting the wires). The rotating head is then directed towards a target, and the camera verifies the incidence of the laser beam. The position of the laser emitted by the rangefinder should be adjusted precisely to hit the target. Similarly, the vertical angle of the laser can be measured by directing the laser to a target at a very shallow angle (S5 Fig). This procedure proves particularly useful for measuring the tilt axis’s angle. Additionally, it is possible to activate the two-axis gimbal and send commands to the motors to execute specific movements, such as 90°, 180°, and 360° rotations. Subsequently, the final laser orientation can be verified against the commanded orientation. This step ensures the alignment between the laser rangefinder and the desired positions dictated by the software.

In situations where altering the placement of the laser rangefinder within the gimbal is not feasible, the laser alignment can be adjusted within the software itself by editing the assumed orientation to match the physical orientation of the laser rangefinder. It is important to note that the physical orientation of the laser emitted by the rangefinder must align with the assumed orientation in the software, not necessarily with the orientation of the rotating box. Thus, the rotating box can be misaligned with the laser as long as the software compensates for this misalignment. However, aligning the laser, rotating box, and software together simplifies the entire process.

### Operation

To make the LiDAR interface simpler and to avoid the need for a LCD screen and more buttons, we decided to define a default initial gimbal orientation, with the user being responsible to align the two-axis gimbal to its default orientation before the start of each scan. The gimbal alignment process is straightforward and consists of: a) levelling the LiDAR scanner on its tripod by using a bubble level attached to the stationary base; b) pointing the rotating head towards the north (bearing = 0°) by using a compass; and c) pointing the laser rangefinder to 40° below the horizon (tilt = −40°), a level in which the laser rangefinder will hit a built-in “tilt stopper” in the 3D-printed case (S6 Fig). The tilt stopper facilitates the positioning of the scanner to the predetermined starting position without the use of additional electronics. An alternative option to the tilt stopper (not used in the current design) is to set the initial gimbal position at the zenith and aligning it with a bubble level before the scan. Finally, since the LiDAR does not have an integrated GPS unit, the scan location must be recorded using an external GPS unit. One simple solution is to use a handheld GPS unit (such as a Garmin GPSMAP 64) placed next to the EcoLiDAR stationary base. The GPS can be used in tracking mode to produce hundreds of waypoints during the LiDAR operation. The points can therefore be averaged to produce a point with higher precision, sometimes with submeter accuracy. Alternatively, it is possible to incorporate a GPS module to the Raspberry Pi Zero to record the GPS location during the scan. Since the EcoLiDAR software code is provided, only minor changes would be required depending on the GPS module of choice.

### Demonstration

To demonstrate the functionality of the LiDAR scanner and the quality of the resultant scan, we mapped a 25 m × 25 m vegetation patch (1.349° N, 103.679° W), located within the National Institute of Education (NIE) campus in Singapore. The site has trees and an irregular soil surface, but no understorey. Field measurements of the tree height are provided in S7 Table. The site was mapped by a single LiDAR scanner, with four scans being performed in a 5 m × 5 m grid, using high resolution settings (1/8 pan axis microstepping and 1/4 tilt-axis microstepping). We used the free software CloudCompare [31] to merge the scans and the statistical environment R [32] with the package lidr [33] to normalize the point cloud and to create some common LiDAR products. We created a digital terrain model, a canopy height model, and a series of figures to visually demonstrate the quality of the scan. For the digital terrain model, we first classified the ground points using a cloth simulation filter algorithm, and created the digital terrain model using a triangular irregular network algorithm [33]. The digital terrain model was subsequently used to normalize the remaining vegetation points, and a canopy height model was then created, also using the triangular irregular network algorithm [33]. An R script with the analysis code is available in the GitHub repository. The height of seven trees within the scanned area were independently measured using a Nikon Coolshot 40i rangefinder and compared with the height estimation from the EcoLiDAR point cloud.

## Results

Since we used a sampling rate of 240 Hz, each scan duration was roughly 95 minutes, with the four scans taking approximately 6 hours and 30 minutes. The LiDAR device performed as expected, producing scans each containing 1,387,200 points at a pan-axis microstepping of 1/8. The resolution of the point cloud is high enough for easy visualization of the main environmental structures, such as trees, branches, and terrain features (Fig 5). The raw scan files as well as the .las version are available at “https://github.com/CalebePMendes/EcoLiDAR.git”.

**Fig 5.**
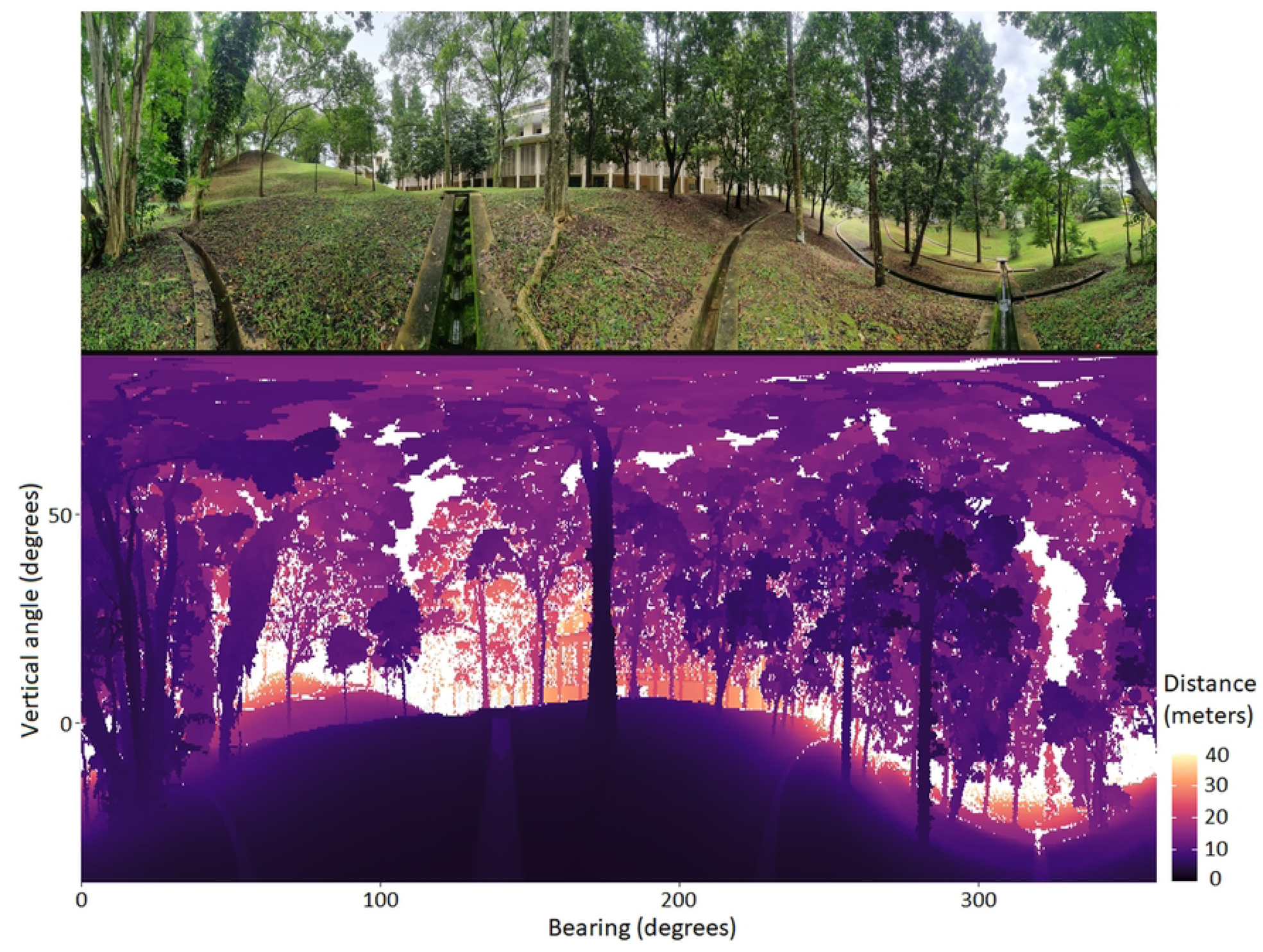
Demonstration site. Panoramic view of the site (top) and the LiDAR product (bottom) from EcoLiDAR using a single scan, with pan axis microstepping of 1/8.

The point cloud obtained from the four scans were successfully aligned and merged, resulting in a single point cloud with 4,954,107 points. The ground points were subsequently identified (Fig 6A) and the digital terrain model was created (Fig 6B). With the normalization of the data using the digital terrain model (Fig 6C), the canopy height model (Fig 7) was obtained. Finally, a 25 m × 25 m area of interest was clipped from the vegetation around to facilitate visualization (Fig 8). The point cloud of a single scan up to the breast height was also displayed to show the cross section of the tree trunks (Fig 9). The tree height estimation based on the point cloud had an average error of −0.24 m (std. dev. = 1 m) when compared to independent measurements using the laser rangefinder (S7 Table).

**Fig 6.**
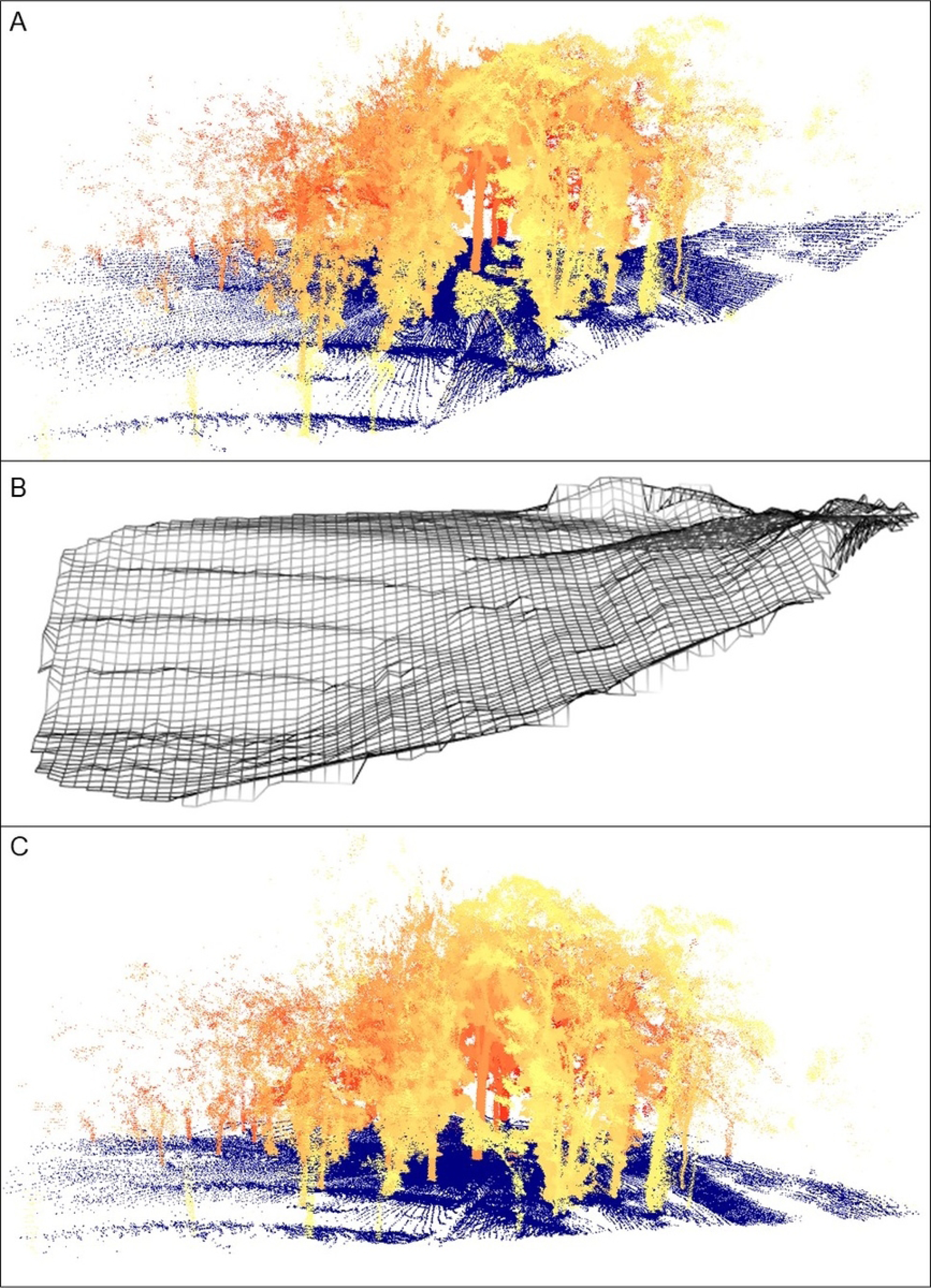
Scan products. Point cloud merged from four scans, with points classified as ground displayed in blue, while the vegetation points are displayed in a yellow-red scale to facilitate depth visualization. A) Raw point cloud, B) the digital terrain model, and C) the resulting normalized point cloud.

**Fig 7.**
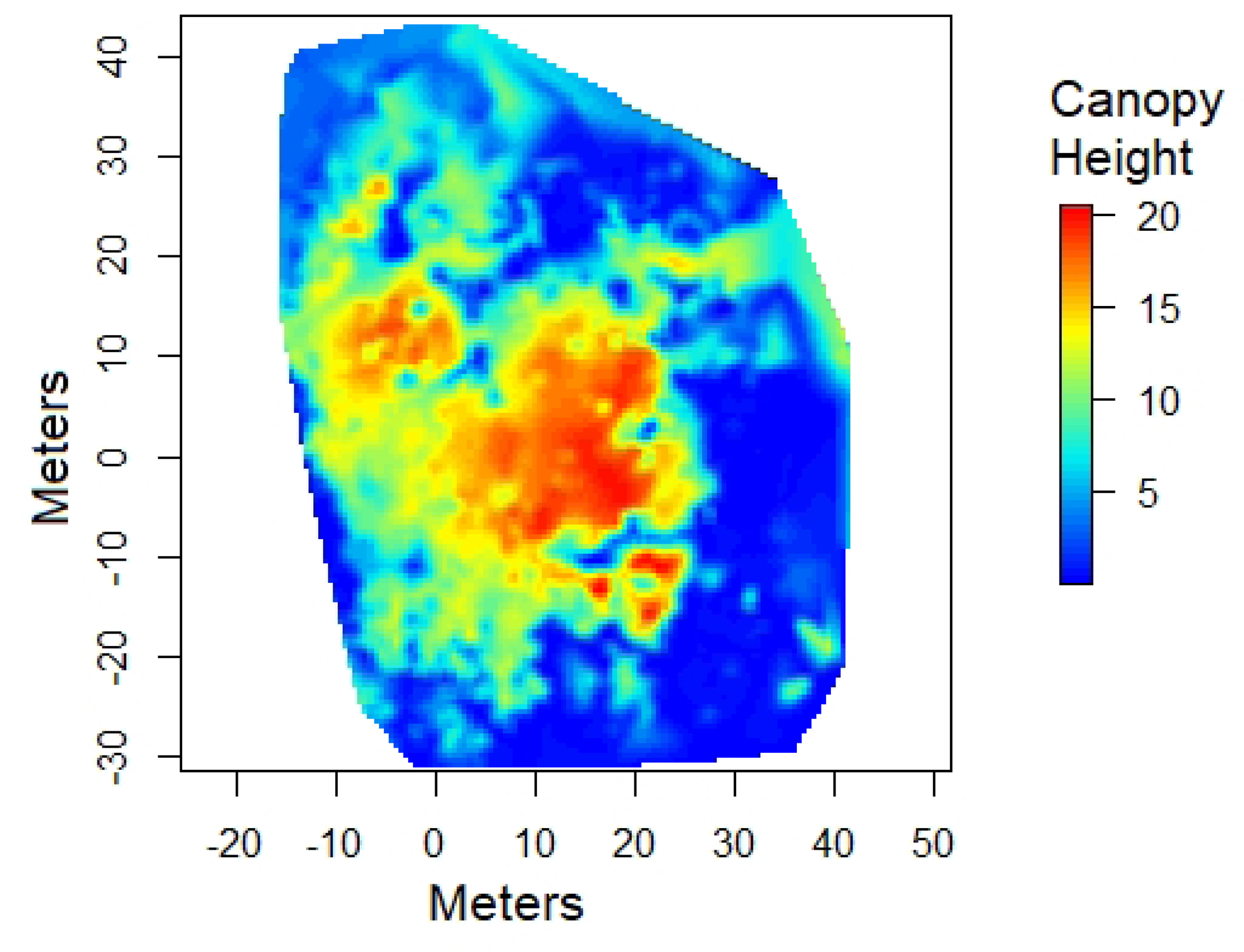
Canopy height model. Canopy height model created from the normalized point cloud (from Fig 6C) using a triangular irregular network algorithm.

**Fig 8.**
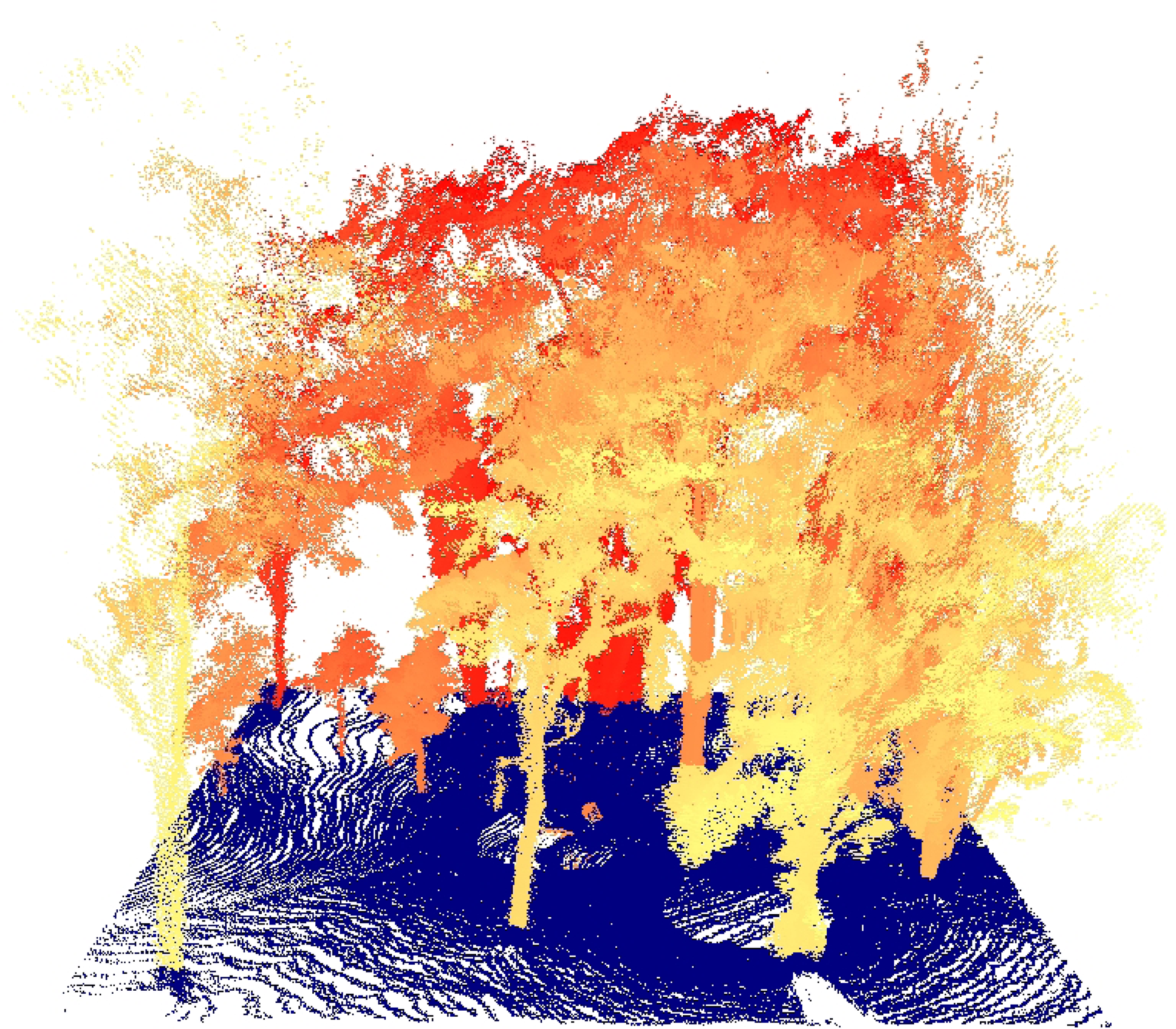
Normalized point cloud. Point cloud of the normalized 25 m × 25 m target area.

**Fig 9.** Single scan point cloud. Point cloud of a single scan showing the profile of trunks at breast height.

In Table 1, we compare the specifications of the EcoLiDAR device with two commercially available “low cost” LiDAR systems. The datasheet of the main components used is also available in the GitHub repository. In terms of price, although we had additional expenses related to development and trial of multiple components, we managed to build a second unit for only 377.89 USD since the design was completed. The price of all components is displayed in S8 Table.

**Table 1.** Performance comparison. A comparison of the specifications of the EcoLiDAR unit with two other “low-cost” commercial LiDAR units.

|  | **EcoLiDAR** | **Faro Focus 120** | **Leica BLK 360** |
| --- | --- | --- | --- |
| Ranging method | Time-of-Flight | Phase-shift | Time-of-Flight |
| Wavelength (nm) | 905 | 905 | 830 |
| Laser Class | Class 1 | Class 3R | Class 1 |
| Beam divergence (mrad) | 8 | 0.19 | 0.4 |
| Ranging accuracy | ±5 cm (at < 2m)<br>±2.5 cm (at > 2m) | ±2 mm | ±4 mm (at 10 m)<br>±7 mm (at 20 m) |
| Max resolution (degrees) | pan: 0.0375, tilt: 0.225<sup>1</sup> | 0.009 | 0.033 |
| Max measurement rate (Hz) | 300 | 976,000 | 360,000 |
| Max range (m) | 40 | 70 | 60 |
| Scanning time | 38 min (at medium<sup>2</sup>)<br>162 min (at high<sup>3</sup>) | 1.5 min | 1.5 min (at medium)<br>3 min (at high) |
| Weight (kg) | 2.2 | 5 | 1 |
| Software | Python script | Faro SCENE | Cyclone REGISTER<br>360 |
| Price (USD) | 400 | <40,000 | 35,000 |

<sup>1</sup> Higher tilt resolutions are possible, but this was the highest we tested.
<sup>2</sup> Medium resolution: 1/8 microstepping on pan axis, 1/4 on tilt axis
<sup>3</sup> High resolution: 1/8 microstepping on pan axis, 1/8 on tilt axis

## Discussion

In this paper, we provide an open-source design to build a functional LiDAR scanner from cheap and widely available electronic components, its potential applications in environmental research, and some limitations. This EcoLiDAR scanner can be built by anyone with limited engineering background, at a fraction of the cost of other “low-cost” commercial units (S8 Table). The device is particularly useful for mapping small areas with static subjects when sub-centimetre precision is not required. Some examples of applications that could benefit from such a device include: mapping the structure of vegetation plots; measuring understorey density at sampling points (such as near camera traps, live traps and dataloggers) within forest patches; producing digital terrain models and canopy height models; and calculating tree breast height diameter and height. Beyond environmental sciences, this LiDAR device can also be used for low-budget engineering and architecture projects, such as for mapping the indoors and outdoors of small buildings, parks, and monuments.

We are fully aware that, performance-wise, our LiDAR system performs poorly when compared with the simplest commercial systems [16] and with other more specialised designs available in the literature [34]. The EcoLiDAR performance and final specification will certainly vary according to the quality of the components used for the build, as well as building techniques used during construction. Therefore, this requires users to perform performance tests for each scanner built before deploying it for data collection. However, we are not aware of any other publication describing a 360° LiDAR designed for maximum simplicity and minimum price possible. By costing a fraction of the cheapest commercial options (∼ 1%; S8 Table), by being far simpler to build than other published designs [34, 35], and by being upgradable according to the needs of each research, our EcoLiDAR scanner opens up a world of new opportunities, particularly for projects in developing countries.

According to the performance requirements of each research project, some simple upgrades can be made to customise the performance of the current LiDAR design. Firstly, since every revolution of the rotating head currently produces a fixed number of measurements, the sampling distribution along the vertical axis of the scan field-of-view is not uniform, with a higher point density towards the azimuth and nadir, and a reduced point density along the horizon. By implementing a software control to change the microstepping according to the tilt angle during the scan, it is possible to correct for this and produce a uniform distribution of points. Such an upgrade would also reduce the scan duration by avoiding oversampling the regions near the azimuth and nadir.

Secondly, the substitution of the LiDAR-Lite v3HP for a laser rangefinder with higher sampling rate can be performed to decrease scan durations. The biggest advantage of having reduced scan durations is to allow for the deployment of larger grids of overlapping scans, which in turn can be merged into a single larger map. A rotating LiDAR sensor, such as the RPLIDAR A1 [36], the Acroname LightWare SF45/B [37] or the Livox Mid-360 [38], are often recommended due to their very high sampling rates. Moreover, since rotating LiDAR sensors scan an entire plane, the current design of two-axis gimbal can be simplified to a simple rotating platform [28]. However, the main drawback of rotating LiDAR sensors are their higher prices and a relative shorter detection range, with the price of 12-m range sensors starting around 100 USD [36] and 50-m range sensors costing around 450 USD [37]. It is noteworthy that low-range LiDAR sensors (∼12 m) can still be useful for indoor applications and for scanning dense understorey vegetation, where a dense grid of close-proximity scans is required in order to overcome the foliage occlusion.

Thirdly, in situations where a high angular resolution is required, some of the possible upgrade options include: a) increasing the pan-axis gear ratio; b) changing the stepper motor drivers for models that allow for finer microstepping; and c) adding a reduction gear to improve the tilt-axis angular resolution. However, as the angular resolution increases, the scan duration increases—a consideration particularly relevant when making extensive 360° scans. A workaround to this issue could be to increase the laser rangefinder sampling rate, which would require the substitution of the LiDAR-Lite v3HP as mentioned above. Another alternative to reduce scan durations is to limit the scan field-of-view to a restricted “scan window” instead of the current 360° panoramic scan. The implementation of a scan window in the current LiDAR design is simple since stepper motors can be equally driven in either clockwise or counter-clockwise directions.

Lastly, in order to detect distant targets, a laser rangefinder capable of reaching the desired distances is required. For such purposes, special attention is required to select a laser with low divergence, in order to keep the laser footprint at long distances as small as possible [1]. The same upgrades described above to improve angular resolutions will also apply to improve the point density for distant targets.

Despite the fact that the EcoLiDAR scanner is capable of providing 3D scans at very low costs, the general low performance still makes its usage inadequate for projects that require higher levels of precision or encompass large areas. Although upgrades can mitigate some of the issues, they also quickly add up in terms of the overall price and complexity, which in turn make the project less attractive to projects with lower budgets. Therefore, it is important to consider alternative options before building this LiDAR scanner. For instance, in situations where a scan range of < 5 m is enough, LiDAR sensors available in modern smartphones and tablets, such as modern iPhones and iPad Pros, can produce excellent results [39, 40]. Similarly, photogrammetry techniques can be used to produce 3D point clouds from photos and videos recorded by regular cameras [41, 42]. Despite the need for a complex post-processing, these photogrammetry methods offer the flexibility of producing high-quality scans from UAV imagery, fulfilling the role of a low-budget substitute to airborne LiDAR.

## Conclusions

To make LiDAR technology more accessible for research in developing countries, we designed a simple ground-based LiDAR scanner that can be assembled under 400 USD (as of 2021) with minimal engineering knowledge. The LiDAR scanner is capable of producing quality 3D point clouds that can be used to create further LiDAR products (e.g., canopy height models, digital terrain models). We also described the device’s limitations, some possible upgrades to improve its performance, and some possible low-cost alternatives. Since the LiDAR scanner is assembled using various components, it is important to evaluate the accuracy and precision of the assembled/modified device before deploying it for data collection. Finally, we hope that the subsystems of this LiDAR design (e.g., two-axis gimbal, laser rangefinder software, stepper motor power settings) may be useful as starting points for better designs in the hands of creative researchers.

## Supporting information captions

**S1 Fig. Pan-axis gear system**. a) stepper motor; b) 20-teeth timing pulley; c) timing belt; d)-60 teeth timing pulley; e) slip ring.

**S2 Code. R code to correct the laser rangefinder offset.**

**S3 Code. Python software to control the LiDAR operation.**

**S4 Manual. User Manual.**

**S5 Fig. Laser alignment process.** The infrared (IR) laser emitted by the rangefinder can be visualized using low-light cameras, webcams or smartphones. This way, the laser orientation in relation to the rotating box can be measured, aligned, and added to the software to produce precise scans. On the left, the IR laser incidence on a rectangular target surface was captured using a video camera. On the right, a line is drawn to point the laser location.

**S6 Fig. Tilt angle stopper.** When the laser rangefinder reaches its starting position, the tilt angle stopper (left) collides with the rotating head’s main box (right).

**S7 Table. Tree height measurements.** Comparison between tree height measurements in field and height estimation based on the EcoLiDAR produced point cloud. All measurements are in meters.

**S8 Table. Price table of the LiDAR components (in USD)**.

